# Glutathione synthetase overexpression in *Acidithiobacillus ferrooxidans* improves halotolerance of iron oxidation

**DOI:** 10.1101/2021.06.29.450459

**Authors:** Yuta Inaba, Alan C. West, Scott Banta

**Author notes:** Corresponding author. Mailing address: Department of Chemical Engineering, Columbia University, 820 Mudd MC4721, 500 W. 120^th^ St., New York, NY 10027.

## Abstract

*Acidithiobacillus ferrooxidans* are well-studied iron- and sulfur-oxidizing acidophilic chemolithoautotrophs that are exploited for their ability to participate in the bioleaching of metal sulfides. Here, we overexpressed the endogenous glutamate--cysteine ligase and glutathione synthetase genes in separate strains and found that glutathione synthetase overexpression increased intracellular glutathione levels. We explored the impact of pH on the halotolerance of iron oxidation in wild type and engineered cultures. The increase in glutathione allowed the modified cells to grow under salt concentrations and pH conditions that are fully inhibitory to wild type cells. These results indicate that glutathione overexpression can be used to increase halotolerance in *A. ferrooxidans* and would likely be a useful strategy on other acidophilic bacteria.

**Importance:** The use of acidophilic bacteria in the hydrometallurgical processing of sulfide ores can enable many benefits including the potential reduction of environmental impacts. The cells involved in bioleaching tend to have limited halotolerance, and increased halotolerance could enable several benefits, including a reduction in the need for fresh water resources. We show that the genetic modification of *A. ferrooxidans* for the overproduction of glutathione is a promising strategy to enable cells to resist the oxidative stress that can occur during growth in the presence of salt.

## Introduction

Acidophilic chemolithotrophic bacteria are critical catalysts in the bioleaching industry as these microorganisms facilitate the solubilization of metals from ores in a less environmentally damaging and energy intensive manner as compared to traditional pyrometallurgical methods (1). In particular, the production of ferric iron plays a significant role in accelerating the dissolution of metal sulfides as it is a strong oxidant (2, 3). Bioleaching technologies have been successfully applied in copper and gold mining; however, many existing mines operate in arid regions, such as Australia or Chile, where water resources are becoming a greater concern (4).

As fresh water becomes more limited for mines, adapting processes can lead to the accumulation of anions in the leach liquor resulting from the intrusion of chloride ions into the water supply from brackish water, dissolution of silicate minerals, and additional recycling of process water (5, 6). The presence of these anions can have a mixed effect on leaching. Small amounts of chloride can have a positive effect on the extraction of copper from the refractory copper sulfide, chalcopyrite (7, 8). The enhancement of copper recovery upon chloride addition is attributed to the reduction of surface passivation of the mineral (9). Furthermore, there are potential new bioleaching applications such as waste metal recycling and corrosion which could benefit from the inclusion of chloride species, permitting unique reaction chemistries (10). However, due to the large transmembrane pH gradient and the positive membrane potential in acidophiles, many anions, other than sulfate, can easily permeate into the cell. The influx of anions causes the cytoplasm to acidify, ultimately causing cell death if the cell is unable to respond to the stress (11). In most species of acidophiles, the toxicity of chloride is exacerbated at increasingly lower pH (12).

Since chloride in known to have a deleterious effect on the better studied acidophiles, some recent efforts have focused on prospecting for new species of acidophilic halophiles which can promote bioleaching under chloride concentrations approaching that of seawater (13-15). The discovery of these new *Acidihalobacter* spp. has spurred the investigation of the genetic mechanisms responsible for their halotolerance (16, 17). It has been shown that these bacterial species have adopted a biphasic salting out approach to increase their internal osmolarity by transporting potassium into the cytoplasm and synthesizing compatible solutes such as proline, ectoine, and periplasmic glucans (18). While the use of these halotolerant species could be promising in future bioleaching processes, their characterization is in early stages and more studies are needed to better understand their suitability for bioleaching. On the other hand, the *Acidithiobacillus* and *Leptospirillum* spp. are widely known, well-characterized acidophiles due to their important role in industrial bioleaching operations (19). These species are represented in mines around the world, but they are limited in their salt tolerance (14). Overall, iron oxidation systems appear to be more sensitive to anions than sulfur oxidation systems, making it difficult to achieve efficient generation of ferric iron when anions are found in leaching liquor (20).

*Acidithiobacillus ferrooxidans* is a key iron- and sulfur-oxidizing bacteria that has well-studied characteristics, available genomic information, and genetic tools and microbiology techniques that have been developed for genetic modifications (21-25). As such, it is becoming an excellent chassis to apply synthetic biology approaches to overcome challenging conditions in novel bioleaching applications (26). As the mechanisms of halotolerance in acidophilic halotolerant species have been uncovered, this offers new opportunities to develop genetically engineered strains of *A. ferrooxidans* that would thrive under even harsher conditions (17). Furthermore, previous studies have reported that genes associated with glutathione in *A. ferrooxidans* respond to various forms of oxidative stress (27-30). And it has been shown that exogenously provided glutathione was effective in reducing intracellular ROS content in an acidophilic microbial consortium containing *A. ferrooxidans*, which suggests that glutathione was an effective ROS scavenger in this system.

Here, we report the investigation of the effects of overexpression of the two genes that form the glutathione biosynthesis pathway in *A. ferrooxidans* in an attempt to improve the halotolerance of the bacteria. By increasing the production of glutathione within the cells, we find that the iron oxidation activity is more resistant to chloride stress at a lower medium pH or higher chloride concentration that is inhibitory to wild type *A. ferrooxidans*. The formation of intracellular ROS species was also measured to better understand how the glutathione enables enhanced iron oxidation in this bacteria.

## Results

### Construction of plasmids and strains overexpressing glutathione biosynthesis genes

The two genes for synthesizing glutathione had been previously identified in the sequenced genome of *A. ferrooxidans* (21). The glutamate--cysteine ligase (*gshA*, AFE_3064) and glutathione synthetase (*gshB*, AFE_3063) sequences were placed behind the *tac* promoter on a pJRD215-based plasmid to generate plasmids pYI43 and pYI46 respectively. These two plasmids were conjugated into *A. ferrooxidans* to successfully generate the AF43 strain expressing *gshA* and the AF46 strain expressing *gshB*.

### Effect of glutathione biosynthesis genes on intracellular glutathione levels

The overexpression of the glutathione biosynthesis genes was expected to increase the production of glutathione in *A. ferrooxidans*. After measuring the extracted concentration of glutathione in the strains, ANOVA showed that AF46 had a significantly higher concentration of glutathione than AF43 or the wild type (WT) (Fig. 1). As AF46 produced more glutathione, we predicted that this strain would be able to better respond to the oxidative stress caused by chloride than wild type stain.

**Fig. 1.**
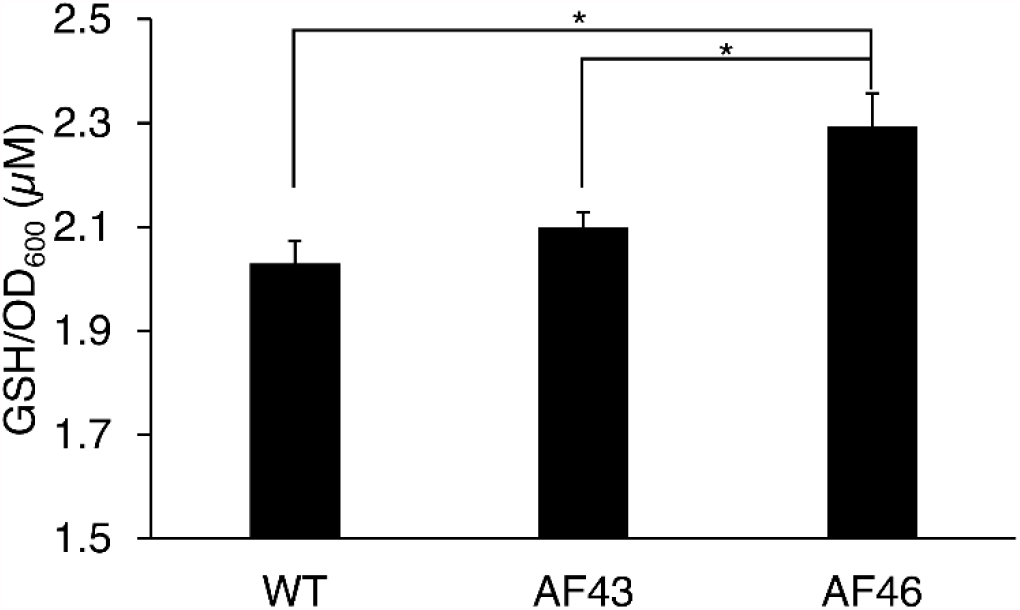
Overexpression of glutathione synthetase, *gshB*, in AF46 increases glutathione concentration compared to WT and AF43. The glutathione concentration from the cell lysates was obtained in triplicate, and error bars indicate standard deviation. Statistical significance was calculated using ANOVA and post-hoc tests (*, p<0.05).

### Preliminary screening of halotolerance

To initially test the effect of the overexpression of increased intracellular glutathione levels in *A. ferrooxidans*, the engineered strains and the WT were grown under varying NaCl concentrations and pH in a 96-well format to determine if enhanced iron oxidation was observed. The assay demonstrated that the halotolerance of *A. ferrooxidans* was sharply impacted within a narrow pH range where iron oxidation could occur as was detected up to 320 mM NaCl at an initial pH of 2.1 and up to 180 mM NaCl at an initial pH of 1.9. However, iron oxidation was not detected beyond 40 mM NaCl at an initial pH of 1.7 (Table S1). Furthermore, while AF46 appeared to have improved tolerance to chloride over WT, AF43 seemed to have no increase in chloride tolerance at all pH conditions. These results suggest that the higher glutathione concentration in the cells resulted in the improved salt tolerance.

### Effect of NaCl concentration and pH on iron oxidation of engineered strains

Since increasing the NaCl concentration or lowering the pH of the media has adverse effects on ferrous iron oxidation by *A. ferrooxidans*, we further investigated each of these effects on WT, AF43, and AF46 cultures. While keeping the NaCl concentration at 50 mM, the effect of the starting pH was compared. As shown in Fig. 2, when the initial pH was lowered incrementally, the strains took longer to complete the conversion of ferrous iron indicating that there was an increasing inhibitory effect of chloride under more acidic conditions. AF46 seemed to oxidize ferrous iron slightly more quickly than WT and AF43 when the starting pH was at 2.04, 1.93, or 1.79. However, when the starting pH was lowered to 1.60, ferrous iron oxidation was only seen with AF46 after a prolonged lag. No oxidation by WT or AF43 was observed even after further incubation, demonstrating that the expression of *gshB* in AF46 extended the pH minimum where iron could be oxidized under chloride stress.

**Fig. 2.**
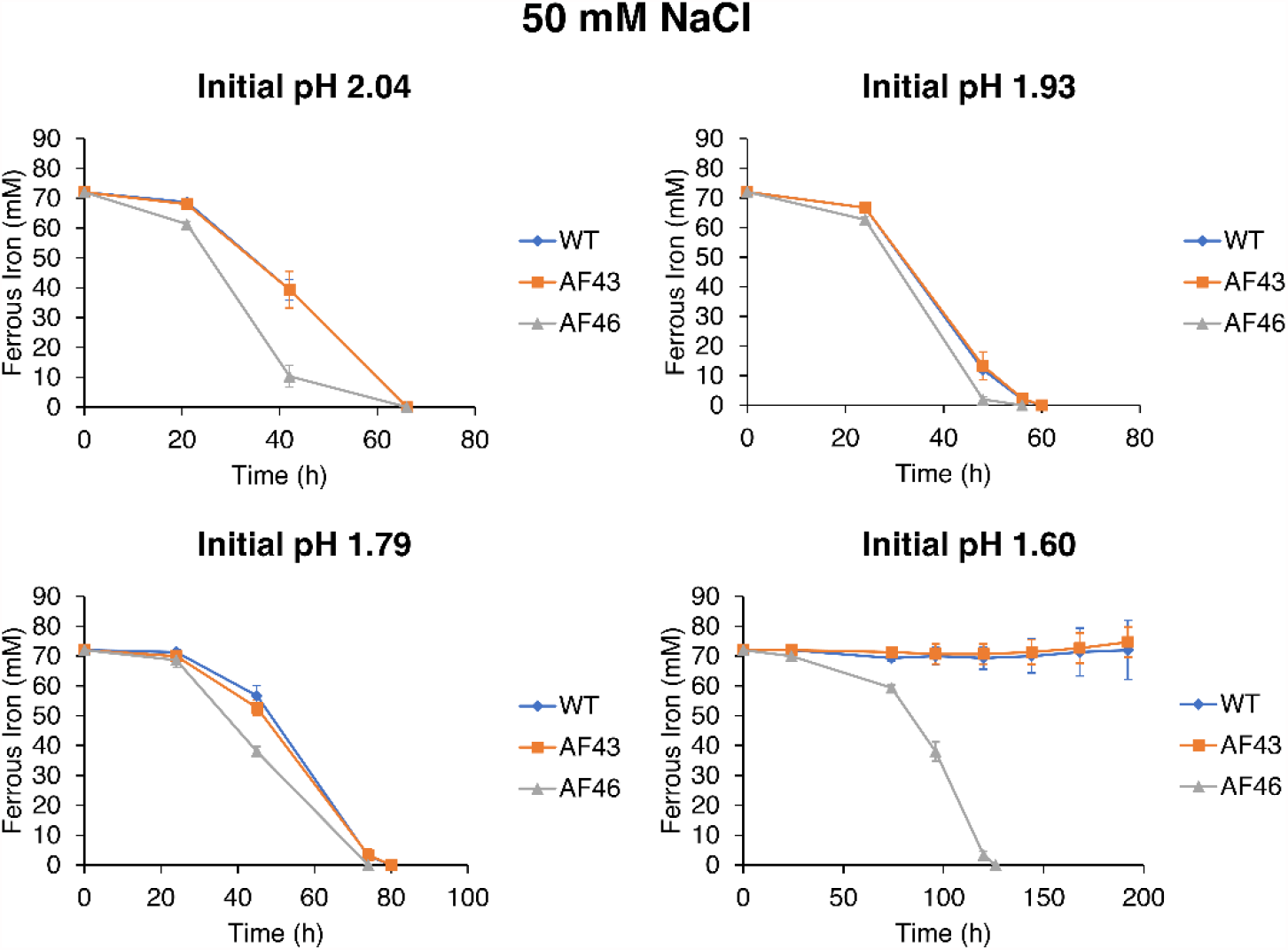
Iron oxidation at constant salt concentration is strongly inhibited when increasingly acidified over a narrow pH range. Iron oxidation activity of WT, AF43, and AF46 was measured in AFM1 medium with 50 mM NaCl at different initial pH. Strains were inoculated into cultures in triplicate, and error bars indicate standard deviation.

We determined that AF46 had higher chloride tolerance at a fixed pH when compared to WT as detected in the preliminary assays. A starting pH of 1.9 was used, which resulted in salt tolerance for iron oxidation in a comparable range to those previously reported in the literature for *A. ferrooxidans* (16, 31). At 150 mM NaCl, the AF46 completed iron oxidation by 144 hours, but only a small amount of iron oxidation was detected with WT, which stagnated at 120 hours. This suggests that this salt concentration was near the MIC for this initial pH (Fig. 3). As all of the ferrous iron was fully oxidized by AF46 under these conditions, a higher NaCl concentration of 200 mM NaCl was also tested. Surprisingly, we found that while AF46 could begin to oxidize iron at this salt level, about 50% of the ferrous iron was oxidized before iron oxidation activity halted (Fig. 3). WT was completely inhibited by 200 mM NaCl. Combined, these results demonstrated that the engineered AF46 strain was capable of iron oxidation under a higher level of chloride stress than WT.

**Fig. 3.**
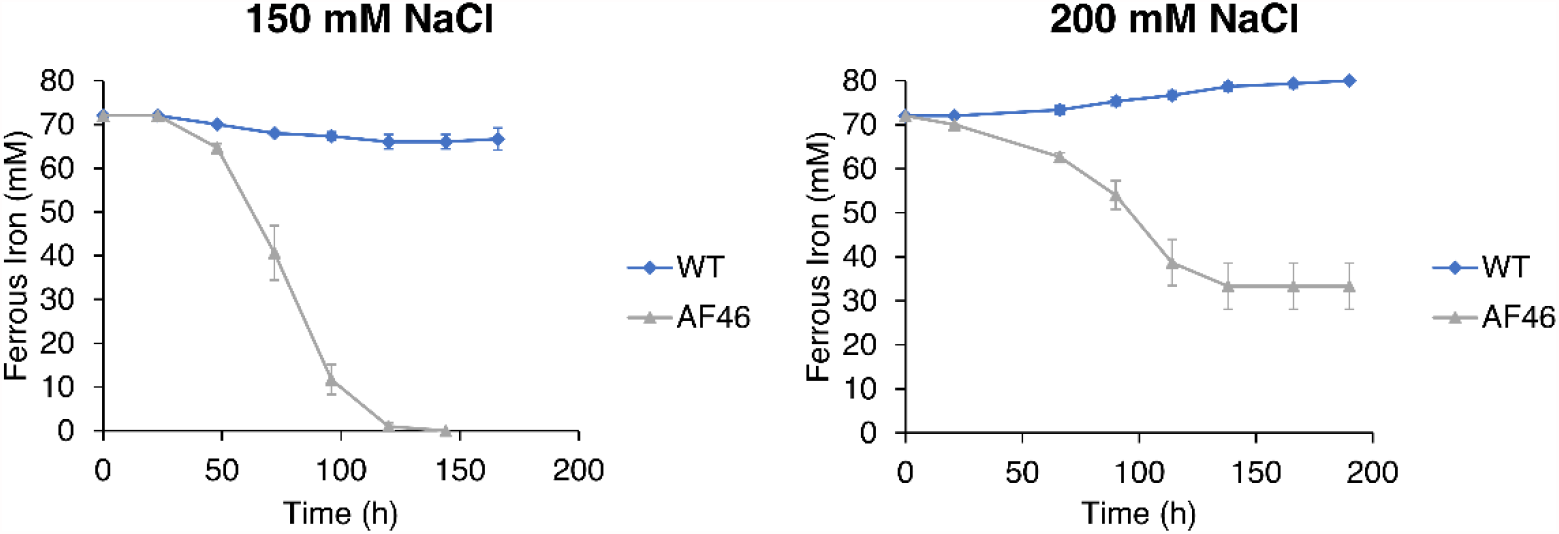
AF46 has increased maximum halotolerance to NaCl compared to WT. Iron oxidation activity of WT and AF46 was measured in AFM1 medium with 150 mM (left) and 200 mM (right) NaCl at an initial pH of 1.9. Strains were inoculated into cultures in triplicate, and error bars indicate standard deviation.

### Impact of glutathione on respiration-induced ROS formation

As increased respiratory activity has been shown to be a response to cytoplasmic acidification in acidophiles, we evaluated whether the differences in iron oxidation halotolerance corresponded to ROS production in the three strains (32). The ROS content in AF46 was significantly higher than WT and AF43 initially (Fig. 4). We found that upon exposure to 100 mM NaCl, the ROS content of all three strains decreased over the first hour of incubation to a similar level. While this salt concentration at the initial pH of 1.9 should be sub-inhibitory for iron oxidation, these results indicate that the initial salt shock induced a similar response. Interestingly, during the entire period of incubation, the ROS content for WT and AF43 remained near this lowered ROS level. While AF46 had a decrease in ROS to a similar level as the two other strains after one hour of incubation, the ROS content increased at 25 hours and approached the ROS levels found in the strain at the start of incubation by 72 hours. The ROS levels in AF46 were significantly higher than WT and AF43 at 25, 45, and 72 hours. These results indicate that the overexpression of *gshB* allows for a higher respiration rate and thus iron oxidation under conditions of oxidative stress induced by chloride in *A. ferrooxidans* after the osmotic shock observed upon initial salt introduction.

**Fig. 4.**
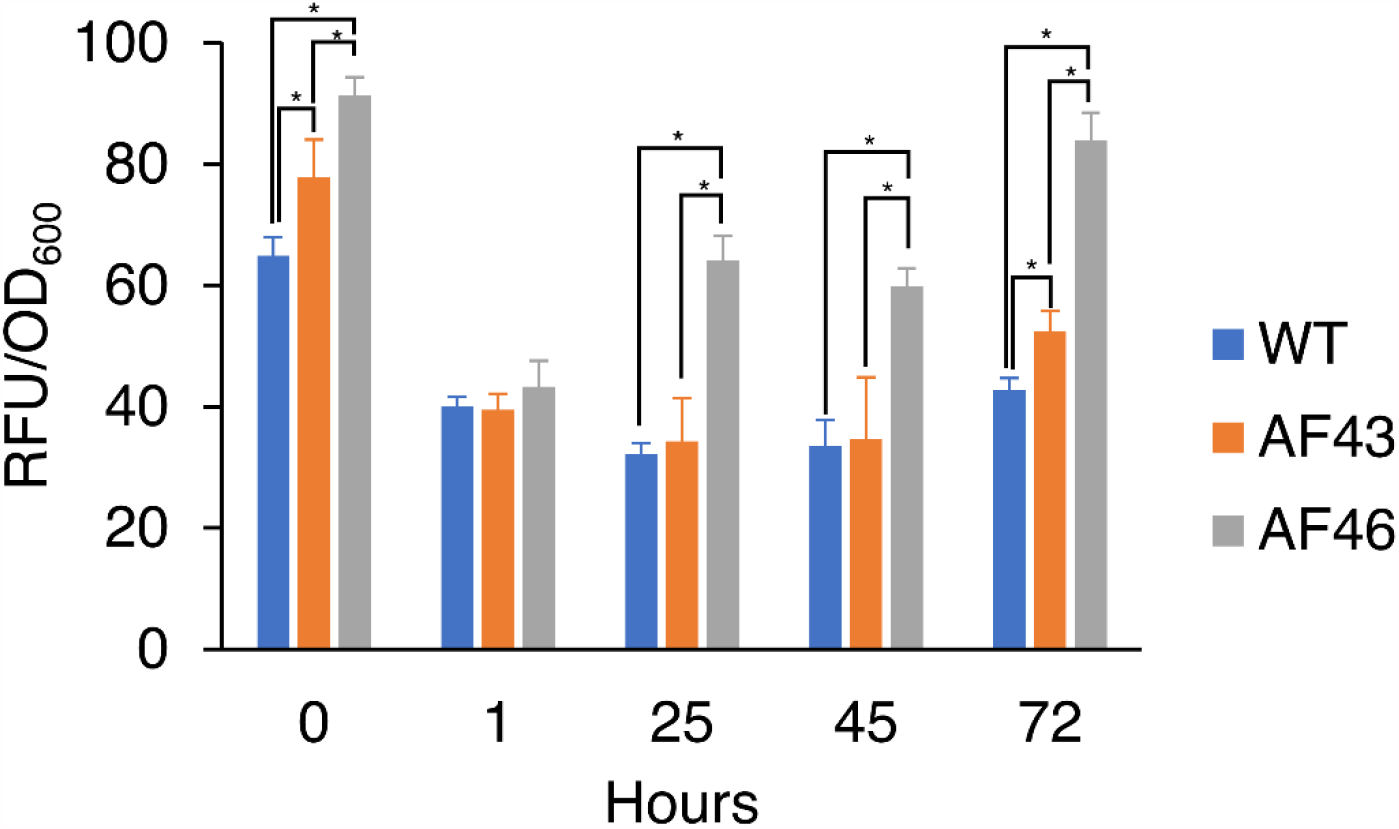
Intracellular ROS levels in WT, AF43, and AF46 exposed to NaCl. Cells were inoculated into AFM1 medium with 100 mM NaCl at an initial pH of 1.9, and ROS content was quantified by measuring the fluorescence of the ROS indicator at various lengths of incubation. Strains were inoculated into cultures in triplicate, and error bars indicate standard deviation. Statistical significance was calculated using ANOVA and post-hoc tests (*, p<0.05).

### Sensitivity to exogenous H_2_O_2_ exposure

Since the glutathione biosynthesis enzymes are localized in the cytoplasm, we evaluated whether AF46 would have an increased tolerance to exogenous sources of oxidative stress where cellular damage would occur starting from the periplasm. High levels of ROS could be generated in acid mine drainage environments, so we treated WT and AF46 with 1 mM H_2_O_2_ for 3 hours where iron oxidation activity had been previously shown to be inhibited with an extended lag phase (29). There was no growth advantage displayed by AF46 compared to the WT, and both strains displayed iron oxidation after 48 hours (Fig. S1). As such, glutathione overexpression does not seem to be effective for all forms of oxidative stress that *A. ferrooxidans* could be exposed to in the environment. This surprising result suggests that *A. ferrooxidans* may utilize different mechanisms of dealing with oxidative stress in the cytoplasm and periplasm.

## Discussion

The iron oxidation activity of *Acidithiobacillus ferrooxidans* has been reported to have low, but varied tolerances to NaCl, which has resulted from the different pH values of the media that have been applied for its growth (5, 14). Microorganisms with iron oxidizing ability tolerant of saline conditions has been noted as a desirable trait in future bioleaching applications due to the importance of ferric iron as an oxidant to solubilize other metals (33). However, many acidophiles have been shown to tolerate lower concentrations of chloride as the medium pH is decreased (12). In this work, we demonstrate the strong impact of medium acidity on the halotolerance of iron oxidation in *A. ferrooxidans* at the low pH values optimal for growth and commonly used to limit ferric iron precipitation. We show that concentrations of chloride that have been indicated previously to be below the MIC for *A. ferrooxidans* can become inhibitory over a narrow pH range, indicating that it may be important to report the halotolerance of acidophiles as a function of pH (16). Further work is needed to better understand if the sharp dependence of halotolerance on pH affects other acidophilic species in a similar fashion.

To address this observation, we have also engineered *A. ferrooxidans* to overexpress glutathione biosynthesis genes, with the goal of augmenting the native response to oxidative stress elicited by NaCl in this species. While acidophilic bacterial species have been reported to have different mechanisms to respond to oxidative stress, *Acidithiobacillus* spp. have a number of unique genes related to glutathione biosynthesis and glutaredoxins (27). These genes involved with glutathione have been reported to be expressed at the early biofilm formation process on pyrite and also protect the cell from oxidative stress induced by cationic heavy metals (30, 34, 35). Although the *gshA* gene has been revealed to be feedback inhibited by glutathione in *E*. coli, we have demonstrated that expression of glutathione synthetase increased intracellular glutathione levels over glutamate--cysteine ligase (36). This suggests that the regulation of these glutathione biosynthesis genes may be different from other studied species as the *gshA* gene from *A. ferrooxidans* was able to sufficiently complement an *E. coli* strain lacking its native gene (37, 38). Although only the overexpression of single genes was tested, simultaneously overexpressing both genes as a single operon may further increase the glutathione concentrations within the cell.

The inhibitory effects of anions has been previously demonstrated to be more pronounced on iron oxidation as compared to sulfur oxidation in *A. ferrooxidans* (20). *A. ferrooxidans* is not known to have genes related to the production of hydroxyectoine or ectoine, which are utilized as osmoprotectants in *Acidihalobacter* and *Leptospirillum* spp., and produces limited amounts of trehalose when exposed to osmotic stress (39). It is possible that producing these osmoprotectants in *A. ferrooxidans* could be an alternative strategy to improving halotolerance. Chloride has been shown to be toxic to acidophiles due to the acidification of the cytoplasm and subsequent ROS generation, and *Leptospirillum ferriphilum* cells respond by increasing respiration rates (32). Because of the limited role of compatible solutes in *A. ferrooxidans*, we had hypothesized that glutathione would be a key compound that would enable the cells to better respond to respiration induced ROS. As predicted, the increased production of glutathione in AF46 allowed for iron oxidation under conditions of lower pH or added chloride concentration compared to WT, confirming that the chloride tolerance was inversely dependent on pH for acidophiles (40). However, we found that ROS levels were decreased when all strains were exposed to NaCl, implying that *A. ferrooxidans* is normally unable to increase respiration rates when faced with chloride stress. These results support a previous proteomics report that a reduction in rusticyanin abundance is observed in *A. ferrooxidans* when grown under high salt conditions and suggests that the additional glutathione content may limit the reduction in the expression of rusticyanin (16).

While increasing glutathione production was effective for enhancing the response of *A. ferrooxidans* to chloride induce oxidative stress, we showed that this did not improve the tolerance of the cells to exogenous hydrogen peroxide which can be generated from surface reactions on metal sulfides (29). Based on our results, overexpressing genes responsible for other molecular mechanisms that have been identified in biofilm cells, such as globin, peroxiredoxin, or alkyl-hydroperoxidase, may allow *A. ferrooxidans* to have increased resistance to bioleaching environments that operate under high pulp densities. The continued investigation of these endogenous proteins and systems under physiologically relevant conditions will be useful in generating other robust strains of *A. ferrooxidans* to use in novel bioleaching operations.

## Materials and Methods

### Chemicals and reagents

All chemicals were sourced from Sigma-Aldrich and enzymes and reagents for DNA manipulation were purchased from NEB unless otherwise noted. A type of dispersed sulfur (#S789400; Toronto Research Chemicals) was used in SM4 medium for the sulfur source as mentioned below (41). All primers used in this study were obtained from Integrated DNA Technologies. Western blot supplies were sourced from Thermo Fisher Scientific.

### Bacterial strains and culturing

*Acidithiobacillus ferrooxidans* ATCC23270 and *Escherichia coli* S17-1 were obtained from ATCC. All *A. ferrooxidan*s cultures were initiated with a starting OD_600_ (optical density measured at 600 nm) of 0.001, corresponding to a cell density of 8.3 × 10^6^ cells/mL. All 100 mL cultures were incubated at 30 °C and shaken at 140 rpm. *A. ferrooxidans* was maintained for use in experiments by weekly subculture into 100 mL of AFM1 medium [0.8 g/L (NH_4_)_2_SO_4_, 0.1 g/L HK_2_PO_4_, 2.0 g/L MgSO_4_·7H_2_O, 5 mL/L trace mineral solution (MD-TMS), and 20.0 g/L FeSO_4_·7H_2_O] at pH 1.8. Cells were harvested by centrifugation at 5,000 x g for 7 min. Harvested cells were kept in 10 mL of AFM1 medium and maintained viability for 1-2 weeks stored at 4 °C.

### Genetic manipulations and confirmation of transformed strains

The pYI11 plasmid was modified to have an in-frame His-tag using the Q5 Site-Directed Mutagenesis with primers pYIHis-F/pYIHis-R as previously described to generate pYI39. This plasmid was then digested with BamHI and KpnI. The *gshA* and *gshB* genes from *A. ferrooxidans* were amplified from genomic DNA prepared using the NucleoSpin Tissue kit (Takara Bio) with primers pYI43-F/pYI43-R and pYI46-F/pYI46-R respectively and purified. The PCR fragments was combined with the digested pYI39 plasmid using NEBuilder HiFi DNA Assembly as per the recommended protocol to generate pYI43 and pYI46. All constructs were transformed into *E. coli* DH5α and were verified by DNA sequencing. DNA sequencing was performed by Genewiz. The primers used for plasmid construction are listed in Table S2. The nucleotide sequences of pYI39, pYI43, and pYI46 have been deposited in Genbank under the accession numbers MZ420731, MZ420732, and MZ420733 respectively.

Plasmids pYI43 and pYI46 were conjugated into *A. ferrooxidans* through the filter mating and conjugation protocol previously described using the donor strain *E. coli* S17-1 with some modifications. After the mating step, possible transconjugant colonies on S204 solid medium plates were grown in AFM1 medium. The *A. ferrooxidans* transconjugants that oxidized iron were then screened in the liquid selection SM4 medium to isolate strains overexpressing *gshA* or *gshB*. After verifying the mutants had persistent kanamycin resistance in SM4 medium, the presence of the overexpressed proteins in the mutants was determined by Western blot. The cell lysates of the transformed *A. ferrooxidans* cells were separated using SDS-PAGE in Novex NuPAGE 4-12% Bis-Tris gel. The separated proteins were transferred onto a Novex 0.45 µm nitrocellulose membrane with a semi-dry blotting unit using the recommended method of the manufacturer. The blotted membrane was blocked using Blocker Fl Fluorescent Blocking Buffer and treated with mouse 6x-His tag monoclonal antibody (HIS.H8) at a 1:1000 dilution. The primary antibody treated membrane was then treated with goat anti-mouse IgG (H+L) cross-adsorbed secondary antibody, Alexa Fluor 488 at a 1:1000 dilution. The fluorescently labeled proteins were detected using a Gel Doc XR+ Imager (Bio-Rad) are shown in Fig. S2 where the expected molecular weight of the His-tagged *gshA* and *gshB* protein are 50.3 and 35.3 kDa respectively. The clone overexpressing *gshA* was referred to as AF43, and the clone overexpressing *gshB* was referred to as AF46. The bacterial strains and plasmids used in this study are listed in Table 1.

**Table 1.** Bacterial strains and plasmids used in this study.

| Strain or Plasmid | Description | Source or Reference |
| --- | --- | --- |
| Strain |  |  |
| *E. coli* S17-1 ATCC 47055 | *recA pro hsdR* RP4-2-Tc::Mu-Km::Tn7 integrated into the chromosome | ATCC |
| *A. ferrooxidans* ATCC 23270 | Type strain | ATCC |
| AF43 | ATCC 23270 with pYI43 | This study |
| AF46 | ATCC 23270 with pYI46 | This study |
| Plasmid |  |  |
| pYI11 | pJRD215 empty vector with *tac* promoter and *rrnB* terminator | (10) |
| pYI39 | pYI11 with in-frame C-terminus His-tag | This study |
| pYI43 | pYI39 with *gshA* from *A. ferrooxidans* | This study |
| pYI46 | pYI39 with *gshB* from *A. ferrooxidans* | This study |

### Preliminary iron oxidation halotolerance assays

Iron oxidation in presence of chloride was conducted in AFM1 medium with the addition of the indicated concentrations of NaCl. The media were acidified with concentrated sulfuric acid to the desired initial pH. Iron oxidation assays investigating the effect of NaCl concentration and initial pH using 96-well plates were incubated at 30 °C without agitation. Iron oxidation by the cells in the wells was noted by the color change of the clear media to an orange-brown indicating that the ferrous iron had been oxidized to ferric iron.

### Cultures evaluating chloride stress and hydrogen peroxide stress on iron oxidation

For investigating the impact of chloride stress on the strains in culture, cells were inoculated into 100 mL of AFM1 medium at the indicated NaCl concentration and initial pH. For investigating the impact of hydrogen peroxide stress, 1 mL of OD_600_ 0.1 cells were washed with AFM1 basal salts [0.8 g/L (NH_4_)_2_SO_4_, 0.1 g/L HK_2_PO_4_, 2.0 g/L MgSO_4_·7H_2_O, and 5 mL/L trace mineral solution (MD-TMS)] at pH 1.8. After washing several times, hydrogen peroxide was added to the cells at a concentration of 1 mM and incubated for 3 hours at 30 °C without agitation. The incubated cells with hydrogen peroxide were directly inoculated into a 100 mL culture of AFM1 medium at pH 1.8. The ferrous iron concentration of cultures was measured by titrating 1 mL of the sampled media mixed with 10 µL of ferroin indicator using a 0.1 M cerium sulfate solution and noting the color change of the solution indicating that the reduced iron species had been oxidized.

### Measurement of intracellular glutathione and intracellular ROS species

To measure intracellular glutathione, WT, AF43, and AF46 cells were grown and harvested cells were washed in AFM1 basal salts then washed in phosphate buffer saline at pH 7.4 and resuspended into a 5% (wt/vol) solution of 5-sulfo-salicylic acid dehydrate. The suspended cells were sonicated to disrupt the cells, and the cell lysate supernatant samples were prepared and assayed as recommended by the manufacturer for use in the Glutathione Colorimetric Detection Kit (Invitrogen). A linear standard for total glutathione concentration from 0 to 6.25 µM was used to obtain glutathione concentrations in the cell lysates from the measured absorbance.

To measure intracellular ROS levels, all strains were initially grown and harvested cells were placed into AFM1 medium with 100 mM NaCl at an initial pH of 1.9. Sampled cells were washed in phosphate buffer saline at pH 7.4, then 2’,7’-dichlorodihydrofluorescein diacetate (H_2_DCFDA) indicator dissolved in dimethyl sulfoxide was added at a final concentration of 10 µM. Samples were incubated in the dark at room temperature for 30 minutes, then the samples were washed with the phosphate buffer saline to remove excess indicator. The fluorescence was measured using an excitation of 495 nm and emission of 522 nm and normalized by the cell density.

### Statistics

All error bars represent one standard deviation from the mean. Statistical analyses using ANOVA and post-hoc tests were conducted using Excel. P-values less than 0.05 were considered to be statistically significant.

## Acknowledgements

This work was supported by the US Army Research Office (W911NF-18-1-0239) and ARPA-E grant DE-AR0001340 from the US Department of Energy. The authors have no competing interests to declare.

## References

1. Johnson DB. 2014. Biomining—biotechnologies for extracting and recovering metals from ores and waste materials. Current Opinion in Biotechnology 30:24–31.

2. Schippers A, Sand W. 1999. Bacterial leaching of metal sulfides proceeds by two indirect mechanisms via thiosulfate or via polysulfides and sulfur. Appl Environ Microbiol 65:319–321.

3. Rohwerder T, Gehrke T, Kinzler K, Sand W. 2003. Bioleaching review part A: progress in bioleaching: fundamentals and mechanisms of bacterial metal sulfide oxidation. Applied Microbiology and Biotechnology 63:239–248.

4. Banerjee I, Burrell B, Reed C, West AC, Banta S. 2017. Metals and minerals as a biotechnology feedstock: engineering biomining microbiology for bioenergy applications. Curr Opin Biotechnol 45:144–155.

5. Zammit CM, Mangold S, rao Jonna V, Mutch LA, Watling HR, Dopson M, Watkin ELJ. 2012. Bioleaching in brackish waters—effect of chloride ions on the acidophile population and proteomes of model species. Applied Microbiology and Biotechnology 93:319–329.

6. Dopson M, Halinen A-K, Rahunen N, Boström D, Sundkvist J-E, Riekkola-Vanhanen M, Kaksonen AH, Puhakka JA. 2008. Silicate mineral dissolution during heap bioleaching. Biotechnology and Bioengineering 99:811–820.

7. Bevilaqua D, Lahti H, Suegama PH, Garcia O, Benedetti AV, Puhakka JA, Tuovinen OH. 2013. Effect of Na-chloride on the bioleaching of a chalcopyrite concentrate in shake flasks and stirred tank bioreactors. Hydrometallurgy 138:1–13.

8. Noguchi H, Okibe N. 2020. The role of bioleaching microorganisms in saline water leaching of chalcopyrite concentrate. Hydrometallurgy 195:105397.

9. Carneiro MFC, Leão VA. 2007. The role of sodium chloride on surface properties of chalcopyrite leached with ferric sulphate. Hydrometallurgy 87:73–82.

10. Inaba Y, West AC, Banta S. 2020. Enhanced microbial corrosion of stainless steel by Acidithiobacillus ferrooxidans through the manipulation of substrate oxidation and overexpression of rus. Biotechnology and Bioengineering 117:3475–3485.

11. Alexander B, Leach S, Ingledew WJ. 1987. The relationship between chemiosmotic parameters and sensitivity to anions and organic acids in the acidophile Thiobacillus ferrooxidans. Microbiology 133:1171–1179.

12. Falagán C, Johnson DB. 2018. The significance of pH in dictating the relative toxicities of chloride and copper to acidophilic bacteria. Research in Microbiology 169:552–557.

13. Huber H, Stetter KO. 1989. Thiobacillus prosperus sp. nov., represents a new group of halotolerant metal-mobilizing bacteria isolated from a marine geothermal field. Archives of Microbiology 151:479–485.

14. Rea SM, McSweeney NJ, Degens BP, Morris C, Siebert HM, Kaksonen AH. 2015. Salt-tolerant microorganisms potentially useful for bioleaching operations where fresh water is scarce. Minerals Engineering 75:126–132.

15. Khaleque HN, Kaksonen AH, Boxall NJ, Watkin ELJ. 2018. Chloride ion tolerance and pyrite bioleaching capabilities of pure and mixed halotolerant, acidophilic iron-and sulfur-oxidizing cultures. Minerals Engineering 120:87–93.

16. Dopson M, Holmes DS, Lazcano M, McCredden TJ, Bryan CG, Mulroney KT, Steuart R, Jackaman C, Watkin ELJ. 2017. Multiple osmotic stress responses in Acidihalobacter prosperus result in tolerance to chloride ions. Frontiers in Microbiology 7.

17. Khaleque HN, González C, Shafique R, Kaksonen AH, Holmes DS, Watkin ELJ. 2019. Uncovering the mechanisms of halotolerance in the extremely acidophilic members of the Acidihalobacter genus through comparative genome analysis. Frontiers in Microbiology 10.

18. Wood JM. 2015. Bacterial responses to osmotic challenges. Journal of General Physiology 145:381–388.

19. Rawlings DE. 2005. Characteristics and adaptability of iron-and sulfur-oxidizing microorganisms used for the recovery of metals from minerals and their concentrates. Microb Cell Fact 4:13.

20. Harahuc L, Lizama HM, Suzuki I. 2000. Selective inhibition of the oxidation of ferrous iron or sulfur in Thiobacillus ferrooxidans. Appl Environ Microbiol 66:1031–1037.

21. Valdes J, Pedroso I, Quatrini R, Dodson RJ, Tettelin H, Blake R, 2nd, Eisen JA, Holmes DS. 2008. Acidithiobacillus ferrooxidans metabolism: from genome sequence to industrial applications. BMC Genomics 9:597.

22. Peng JB, Yan WM, Bao XZ. 1994. Plasmid and transposon transfer to Thiobacillus ferrooxidans. J Bacteriol 176:2892–7.

23. Wang H, Liu X, Liu S, Yu Y, Lin J, Lin J, Pang X, Zhao J. 2012. Development of a markerless gene replacement system for Acidithiobacillus ferrooxidans and construction of a pfkB mutant. Applied and Environmental Microbiology 78:1826–35.

24. Kernan T, Majumdar S, Li X, Guan J, West AC, Banta S. 2016. Engineering the iron-oxidizing chemolithoautotroph Acidithiobacillus ferrooxidans for biochemical production. Biotechnology and Bioengineering 113:189–97.

25. Inaba Y, Banerjee I, Kernan T, Banta S. 2018. Transposase-mediated chromosomal integration of exogenous genes in Acidithiobacillus ferrooxidans. Applied and Environmental Microbiology 84:e01381–18.

26. Gumulya Y, Boxall N, Khaleque H, Santala V, Carlson R, Kaksonen A. 2018. In a quest for engineering acidophiles for biomining applications: challenges and opportunities. Genes 9:116.

27. Cárdenas JP, Moya F, Covarrubias P, Shmaryahu A, Levicán G, Holmes DS, Quatrini R. 2012. Comparative genomics of the oxidative stress response in bioleaching microorganisms. Hydrometallurgy 127-128:162–167.

28. Orell A, Navarro CA, Arancibia R, Mobarec JC, Jerez CA. 2010. Life in blue: Copper resistance mechanisms of bacteria and Archaea used in industrial biomining of minerals. Biotechnology Advances 28:839–848.

29. Bellenberg S, Huynh D, Poetsch A, Sand W, Vera M. 2019. Proteomics reveal enhanced oxidative stress responses and metabolic adaptation in Acidithiobacillus ferrooxidans biofilm cells on pyrite. Frontiers in Microbiology 10.

30. Xia J-l, Wu S, Zhang R-y, Zhang C-g, He H, Jiang H-c, Nie Z-y, Qiu G-z. 2011. Effects of copper exposure on expression of glutathione-related genes in Acidithiobacillus ferrooxidans. Current Microbiology 62:1460–1466.

31. Rivera-Araya J, Huynh ND, Kaszuba M, Chávez R, Schlömann M, Levicán G. 2020. Mechanisms of NaCl-tolerance in acidophilic iron-oxidizing bacteria and archaea: Comparative genomic predictions and insights. Hydrometallurgy 194:105334.

32. Rivera-Araya J, Pollender A, Huynh D, Schlömann M, Chávez R, Levicán G. 2019. Osmotic imbalance, cytoplasm acidification and oxidative stress induction support the high toxicity of chloride in acidophilic bacteria. Frontiers in Microbiology 10.

33. Kaksonen AH, Deng X, Bohu T, Zea L, Khaleque HN, Gumulya Y, Boxall NJ, Morris C, Cheng KY. 2020. Prospective directions for biohydrometallurgy. Hydrometallurgy 195:105376.

34. Vera M, Krok B, Bellenberg S, Sand W, Poetsch A. 2013. Shotgun proteomics study of early biofilm formation process of Acidithiobacillus ferrooxidans ATCC 23270 on pyrite. Proteomics 13:1133–1144.

35. Zheng C, Zhang L, Chen M, Zhao XQ, Duan Y, Meng Y, Zhang X, Shen RF. 2018. Effects of cadmium exposure on expression of glutathione synthetase system genes in Acidithiobacillus ferrooxidans. Extremophiles 22:895–902.

36. Murata K, Kimura A. 1990. Overproduction of glutathione and its derivatives by genetically engineered microbial cells. Biotechnology Advances 8:59–96.

37. Ohtake Y, Watanabe K, Tezuka H, Ogata T, Yabuuchi S, Murata K, Kimura A. 1989. Expression of the glutathione synthetase gene of Escherichia coli B in Saccharomyces cerevisiae. Journal of Fermentation and Bioengineering 68:390–394.

38. Powles R, Deane S, Rawlings D. 1996. The gene for γ-glutamylcysteine synthetase from Thiobacillus ferrooxidans has low homology to its Escherichia coli equivalent and is linked to the gene for citrate synthase. Microbiology 142:2543–2548.

39. Galleguillos PA, Grail BM, Hallberg KB, Demergasso CS, Johnson DB. 2018. Identification of trehalose as a compatible solute in different species of acidophilic bacteria. Journal of Microbiology 56:727–733.

40. Suzuki I, Lee D, Mackay B, Harahuc L, Oh JK. 1999. Effect of various ions, pH, and osmotic pressure on oxidation of elemental sulfur by Thiobacillus thiooxidans. Applied and environmental microbiology 65:5163–5168.

41. Inaba Y, Kernan T, West AC, Banta S. 2021. Dispersion of sulfur creates a valuable new growth medium formulation that enables earlier sulfur oxidation in relation to iron oxidation in Acidithiobacillus ferrooxidans cultures. Biotechnology and Bioengineering Advance online publication.

